# Electrophysiological response geometries reveal shared and protocol-specific cell-type organization

**DOI:** 10.64898/2026.01.08.698522

**Authors:** Zihan Yang, Yuchen Xiao

## Abstract

Neuronal electrophysiological identity is expressed through responses to input, but how this identity is organized across distinct physiological input regimes remains unclear. Conventional descriptors such as firing rate, threshold, latency, or response amplitude capture local aspects of excitability, while different current-clamp protocols expose different dimensions of cellular response. Here, we asked whether protocol-specific response measurements reveal distinct physiological axes while preserving shared organization among genetically targeted cell groups. Using intracellular current-clamp recordings from mouse visual cortex, we constructed multivariate response geometries from long-pulse, ramp, and brief-pulse stimulation protocols, respectively probing sustained spiking output, dynamic recruitment, and fast transient response/gain. Each protocol revealed a structured and interpretable geometry: long-pulse responses organized cells along sustained-output and high-current response dimensions, ramp responses isolated recruitment-threshold structure, and brief-pulse responses captured fast transient response and gain-related variation. Driver-defined groups occupied ordered positions within these geometries, indicating that protocol-specific response axes were linked to cell-group organization. Across protocols, driver-defined group distance structures remained positively aligned after balanced resampling and comparison with driver-label shuffle null models, showing that protocol-specific geometries retained partially shared relational structure. These findings support a protocol-resolved view of cellular electrophysiological identity as a multidimensional response organization, in which distinct physiological probes reveal protocol-specific response modes embedded within shared cell-group structure.

## Introduction

Neurons transform input into electrical responses through coordinated interactions among membrane excitability, spike initiation, adaptation, and intrinsic conductance dynamics. These responses vary widely across neuronal populations, but this variation is structured, reflecting the diversity of cortical cell classes and their characteristic cellular response properties^1–3^. Some cells require stronger drive to enter the spiking regime, some sustain high-frequency output under maintained depolarization, and others respond more strongly to brief or subthreshold inputs. A central challenge is therefore to understand how these response properties are organized across cell types, rather than treating them as independent electrophysiological measurements.

Cell-type-related electrophysiological differences are often described using scalar features such as firing rate, rheobase, spike threshold, latency, adaptation, input resistance, or voltage-response amplitude. These measurements are essential, but each captures only one aspect of cellular excitability. Intrinsic electrophysiological identity may instead be better represented as a multivariate organization: a response space in which cells occupy structured positions, major axes reflect interpretable physiological variation, and cell groups can be compared through their relative locations and distances^2,4,5^.

This organization depends on the stimulation protocol used to probe the cell, as standardized current-clamp recordings can expose different intrinsic response properties across stimulus regimes^2,4^. Different current-clamp protocols impose different temporal and dynamical demands on intrinsic physiology. Long-pulse stimulation emphasizes sustained spiking output under maintained depolarization; ramp stimulation emphasizes dynamic recruitment and threshold demand during gradually increasing input; and brief-pulse stimulation emphasizes fast transient voltage response and input-normalized gain. These protocols therefore provide complementary windows onto cellular excitability. The central question of this study is whether protocol-specific response geometries reveal distinct physiological dimensions while preserving shared organization among driver-defined cell groups.

To address this question, we analyzed intracellular current-clamp recordings from genetically targeted mouse visual cortical neurons from the Allen Institute Patch-seq resource^4,6^. For each protocol, we extracted protocol-native response features, aggregated sweep-level measurements into cell-level representations, and constructed low-dimensional response geometries. We then quantified driver-defined cell-group organization within and across protocols using principal component embeddings, feature loadings, centroid positions, pairwise distance structures, balanced resampling, and driver-label shuffle controls.

We find that intrinsic electrophysiological organization emerges within and across stimulation regimes. Within individual protocols, cells are organized along interpretable physiological axes: long-pulse responses define a sustained spiking-output geometry, ramp responses isolate a dynamic recruitment geometry, and brief-pulse responses reveal fast transient response and gain-related structure. Across protocols, these geometries retain partially preserved driver-group relational structure, indicating that different stimulation regimes reveal distinct physiological projections of a shared intrinsic organization. Together, these results establish protocol-resolved response geometry as a framework for connecting multivariate single-cell physiology to biological cell-type organization.

## Results

### From stimulation protocols to electrophysiological response geometries

Intrinsic electrophysiological identity cannot be fully captured by a single response measurement or a single stimulation protocol. A neuron that appears similar to another under sustained depolarization may differ in the current required for spike recruitment, and both may differ again in their sensitivity to brief transient input. This raises the central question addressed here: whether genetically targeted single neurons, grouped by driver line, are organized by a shared intrinsic electrophysiological structure that persists across stimulation regimes, or whether each protocol defines a largely independent response organization ^2,4,5,7^.

To address this question, we built a protocol-resolved framework using intracellular current-clamp recordings from genetically labeled mouse visual cortical neurons in the Allen Institute Patch-seq dataset^8,9^**(Fig. 1A-B)**. We analyzed three stimulation protocols that impose distinct input demands on the cell. Long-pulse stimulation measures sustained spiking output under maintained current input, ramp stimulation measures the current and timing required for dynamic spike recruitment, and brief-pulse stimulation measures fast transient voltage responses and input-normalized gain. These protocols therefore provide complementary views of intrinsic excitability: sustained output, recruitment threshold, and transient input sensitivity.

**Figure 1.**
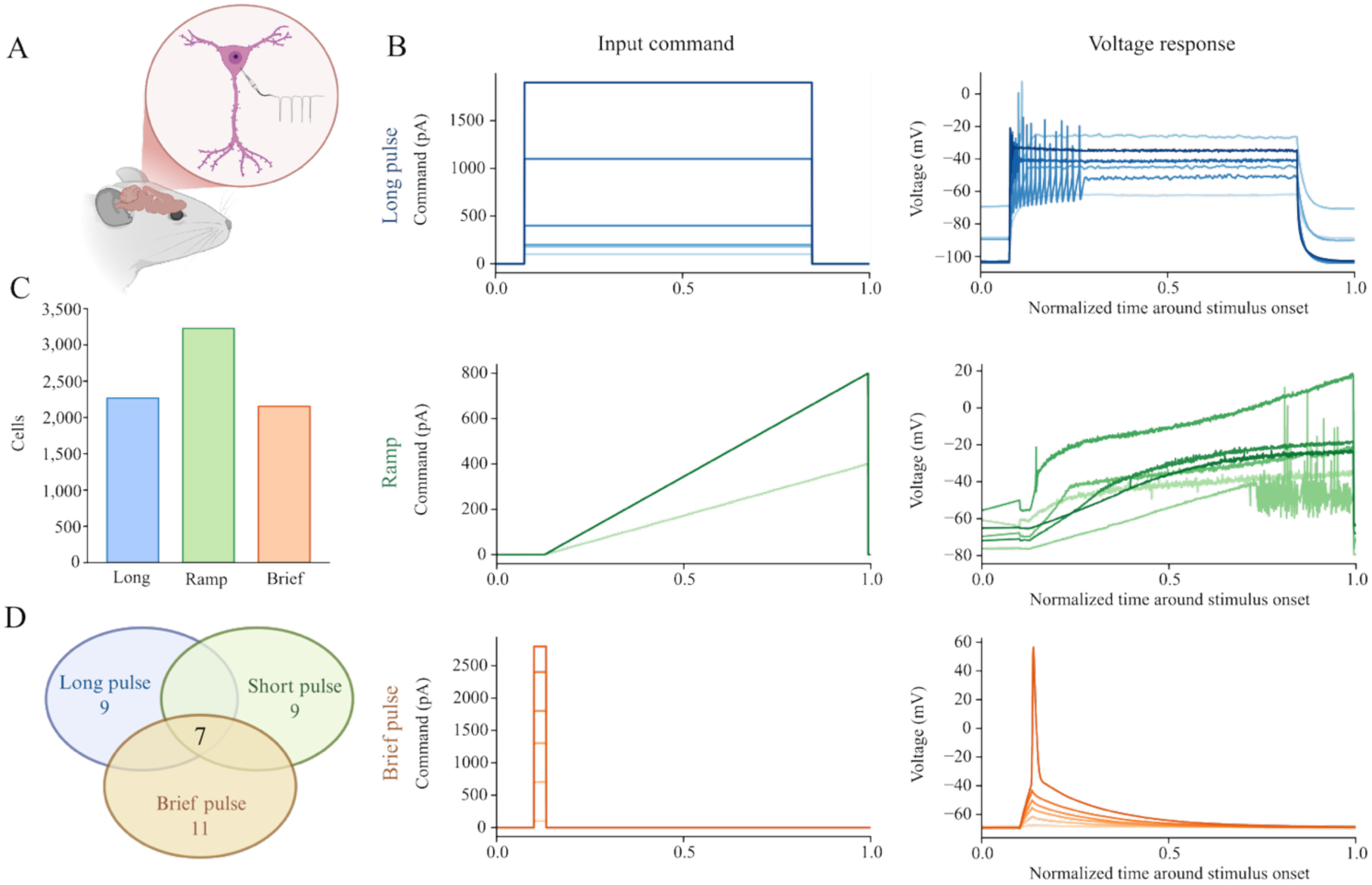
Protocol-resolved construction of electrophysiological response geometries. **(A)** Schematic overview of intracellular current-clamp recordings from genetically labeled mouse visual cortical neurons. **(B)** Representative current commands and voltage responses for long-pulse, ramp, and brief-pulse stimulation. Long-pulse stimulation was used to define a sustained spiking-output geometry, ramp stimulation a dynamic recruitment geometry, and brief-pulse stimulation a fast transient response/gain geometry. Trace intensity denotes increasing input amplitude or stimulation strength; time is normalized within each protocol for visualization. **(C)** Number of cells included in PCA-based response geometry construction for each stimulation protocol. **(D)** Driver-line coverage across protocol-specific geometries, including the driver-defined groups shared across protocols and used for balanced cross-protocol validation.

We converted each protocol into a cell-level response geometry by transforming sweep-level recordings into protocol-native electrophysiological features. A shared quality-control framework retained sweeps with valid current-clamp traces, stable baseline activity, identifiable stimulus structure, and measurable voltage responses, with additional protocol-specific checks matched to long-pulse, ramp, or brief-pulse waveform structure. QC-passed sweeps were aggregated into one feature vector per cell for each protocol, producing separate long-pulse, ramp, and brief-pulse feature tables for geometry construction **(Fig. 1C-D**; **Table 1)**.

**Table 1:**
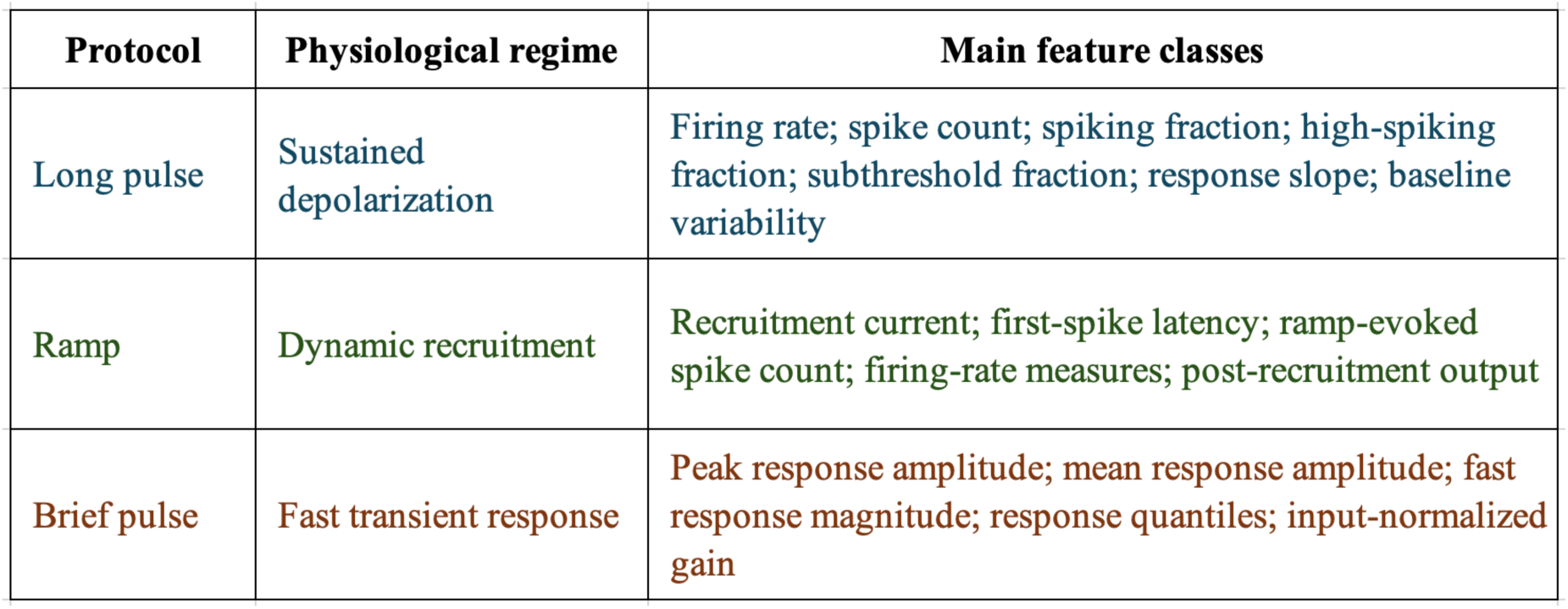
Protocol-specific feature classes. Long-pulse, ramp, and brief-pulse stimulation probe sustained depolarization, dynamic recruitment, and fast transient responses, respectively. Cell-level response geometries were constructed from protocol-native feature classes matched to each physiological regime.

| Protocol | Physiological regime | Main feature classes |
| --- | --- | --- |
| Long pulse | Sustained depolarization | Firing rate; spike count; spiking fraction; high-spiking fraction; subthreshold fraction; response slope; baseline variability |
| Ramp | Dynamic recruitment | Recruitment current; first-spike latency; ramp-evoked spike count; firing-rate measures; post-recruitment output |
| Brief pulse | Fast transient response | Peak response amplitude; mean response amplitude; fast response magnitude; response quantiles; input-normalized gain |

Each protocol-specific feature table was treated as a multivariate electrophysiological response space rather than a collection of isolated scalar measurements. After feature standardization, cells were projected into principal component spaces, allowing major axes of response variation to be interpreted through their physiological feature loadings. Within each protocol, driver-line centroids were computed in PC space, and pairwise Euclidean distances between centroids were used to quantify relational organization among driver-defined cell groups. This produced two linked representations for each stimulation regime: a cell-level response geometry and a driver-line relational geometry.

This framework establishes the logic of the following analyses. First, we test whether each stimulation protocol defines a coherent and physiologically interpretable response geometry. Second, we identify which driver-defined groups occupy the high and low ends of the major response axes, linking multivariate geometry to biological cell-group structure. Third, we test whether the relational organization among driver-defined groups is preserved across stimulation regimes. In this way, long-pulse, ramp, and brief-pulse recordings are treated not as redundant assays, but as complementary probes for decomposing intrinsic electrophysiological identity into shared and protocol-specific response dimensions.

### Long-pulse stimulation defines a sustained-output response geometry

Long-pulse stimulation directly tests how single neurons transform maintained depolarizing input into sustained spiking output, a core component of standardized intrinsic electrophysiological phenotyping^2^. We therefore used long-pulse-derived features to determine whether maintained current injection defines an organized electrophysiological response geometry across genetically targeted cells. Each cell was represented by its long-pulse response profile and projected into principal component space.

The resulting feature space was low-dimensional and structured. A small number of principal components captured most of the variance in long-pulse responses **(Fig. 2A)**, and the PC1–PC2 embedding revealed a continuous response geometry rather than discrete isolated clusters **(Fig. 2B)**. Driver-defined groups were not randomly distributed within this continuum. Instead, they occupied biased regions of the long-pulse space, indicating that sustained stimulation captures coordinated response variation linked to cell-group identity without forcing genetically targeted cells into sharply separated categorical clusters.

**Figure 2.**
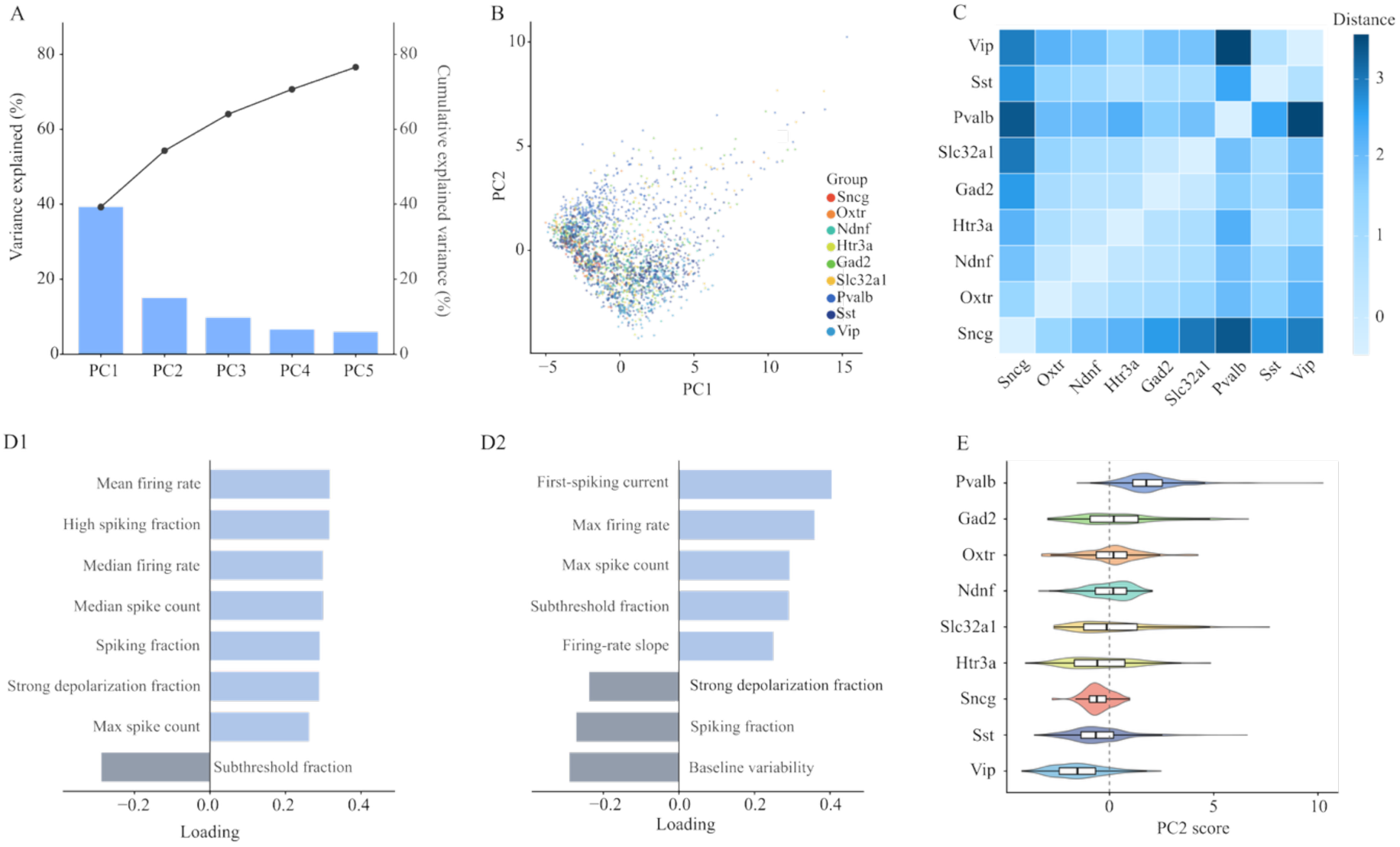
Long-pulse stimulation defines a sustained spiking-output geometry. **(A)** Variance explained by principal components derived from long-pulse cell-level response features. Bars indicate variance explained by individual PCs, and the line indicates cumulative explained variance. **(B)** Cell-level PC1–PC2 embedding of long-pulse response features, with cells colored by driver-defined group. Driver-line labels are abbreviated by primary marker genes for readability. **(C)** Pairwise Euclidean distances between driver-line centroids in long-pulse PC space. **(D)** Top feature loadings for PC1 and PC2. PC1 primarily reflected sustained spiking-output strength and the subthreshold-to-spiking transition, whereas PC2 captured recruitment and high-current output-related variation. **(E)** Driver-defined group distributions along PC2, showing systematic shifts across genetically targeted cell groups.

Feature loadings identified the physiological dimensions underlying this geometry. PC1 was dominated by sustained spiking-output features, including firing-rate measures, spike-count measures, high-spiking fraction, and spiking fraction, with subthreshold fraction loading in the opposite direction **(Fig. 2D1)**. PC1 therefore captured the major transition from weak or subthreshold responses to robust sustained spiking output. PC2 captured a complementary output-recruitment dimension, with positive loadings from first-spiking current, maximum firing rate, maximum spike count, subthreshold fraction, and firing-rate slope, and negative loadings from features including spiking fraction and baseline variability **(Fig. 2D2)**. Thus, long-pulse stimulation did not simply sort cells by average firing rate; it organized sustained output, recruitment demand, and high-current response intensity into a multivariate physiological geometry.

This geometry also revealed ordered driver-line structure. Driver-line centroids were separated in PC space **(Fig. 2C)**, and PC2 distributions identified the groups occupying the extremes of the output-recruitment axis **(Fig. 2E)**. Pvalb-targeted cells were shifted toward the higher end of PC2, consistent with the fast-spiking and high-frequency output properties commonly associated with Pvalb interneurons^1,10^, whereas Vip-targeted cells occupied the lower end of the distribution. Other driver-defined groups occupied intermediate positions, indicating that long-pulse stimulation organizes genetically targeted cells along a graded sustained output-recruitment continuum rather than a binary division between cell classes. Together, these results establish long-pulse stimulation as a sustained-output assay that links multivariate intrinsic response features to structured driver-defined cell-group organization.

### Ramp stimulation reveals a recruitment-threshold response geometry

Ramp stimulation tests a different input-output regime from sustained current injection. Because the command current increases gradually over time, this protocol directly measures when a cell enters the spiking regime and how much input is required for recruitment. We therefore used ramp-derived features to construct a dynamic recruitment geometry, capturing recruitment current, first-spike latency, spike count, and firing-rate responses during gradually increasing input.

The ramp feature space formed a structured response geometry with driver-line-dependent occupancy **(Fig. 3A-B)**. As in the long-pulse geometry, cells did not form isolated driver-line clusters. Instead, driver-defined groups occupied biased regions within a shared response continuum. This establishes ramp stimulation as an organized cell-group assay, but one centered on a different physiological dimension: not sustained output under maintained drive, but the threshold and timing of spike recruitment during increasing input.

**Figure 3.**
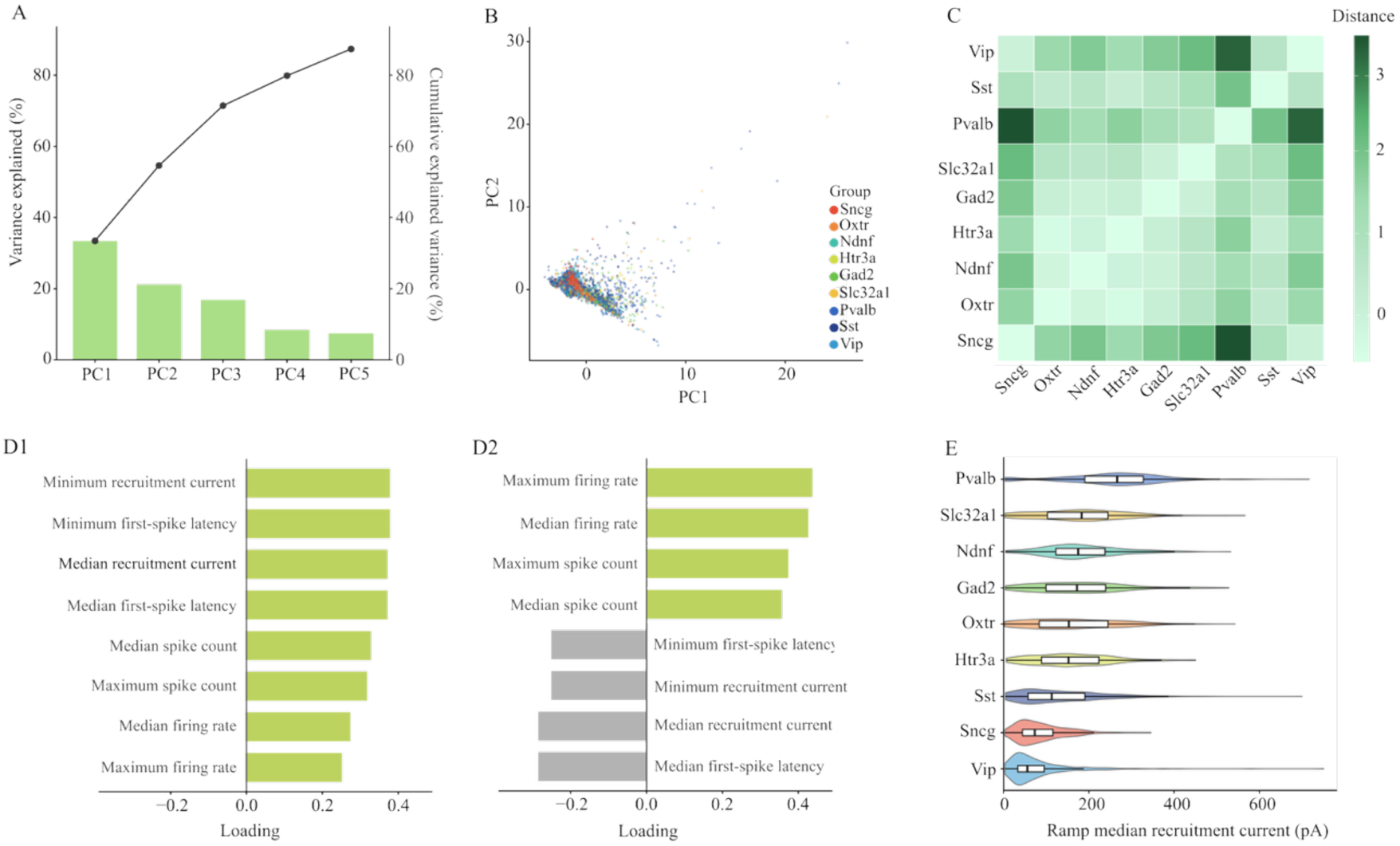
Ramp stimulation isolates a dynamic recruitment geometry. **(A)** Variance explained by principal components derived from ramp-response features. Bars indicate variance explained by individual PCs, and the line indicates cumulative explained variance. **(B)** Cell-level PC1–PC2 embedding of ramp-response features, with cells colored by driver-defined population. Driver-line names are abbreviated by primary marker genes for readability. **(C)** Pairwise Euclidean distances between driver-line centroids in ramp PC space. **(D)** Top feature loadings for PC1 and PC2. PC1 was dominated by recruitment-current and first-spike latency features, whereas PC2 reflected firing-rate and spike-count output during ramp stimulation. **(E)** Driver-defined population distributions of median recruitment current, providing a raw physiological anchor for the ramp recruitment axis.

Feature loadings confirmed that recruitment demand was the dominant organizing principle of the ramp geometry. PC1 was strongly defined by recruitment-current and first-spike-latency features, including minimum recruitment current, median recruitment current, and latency to first spike **(Fig. 3D1)**. Cells with higher PC1 scores required stronger or later ramp input before spiking, whereas cells with lower PC1 scores were recruited under weaker ramp drive. PC2 captured complementary output-related variation, with prominent contributions from firing-rate and spike-count features once spiking had been recruited **(Fig. 3D2)**. Thus, ramp stimulation separated two linked but distinct physiological processes: the input required to initiate spiking and the output produced after recruitment.

Driver-defined groups were ordered along this recruitment axis. Driver-line centroids showed structured pairwise distances in ramp PC space **(Fig. 3C)**, and the raw median recruitment-current distributions provided a direct physiological anchor for this organization **(Fig. 3E)**. Pvalb-targeted cells were shifted toward higher recruitment currents^2,3^, whereas Vip- and Sncg-targeted cells occupied the lower-current end of the distribution, with other driver-defined groups positioned between these extremes. This ordering demonstrates that ramp stimulation organizes genetically targeted cells along a graded recruitment-threshold continuum.

Together, these results establish ramp stimulation as a dynamic recruitment assay that complements and extends the sustained-output geometry defined by long-pulse stimulation. Whereas long-pulse responses organize cells by sustained spiking output and high-current response structure, ramp responses isolate the current and timing required to enter the spiking regime. The high-recruitment-current position of Pvalb-targeted cells and the low-recruitment-current position of Vip-targeted cells parallel their positions in the long-pulse geometry, supporting a convergent output-threshold organization across these two stimulation regimes.

### Brief-pulse stimulation reveals a transient-gain response geometry

Brief-pulse stimulation probes a response regime that is not captured by either sustained depolarization or gradual ramp recruitment. Whereas long-pulse stimulation emphasizes maintained spiking output and ramp stimulation emphasizes the current required to enter the spiking regime, brief-pulse stimulation measures the immediate voltage response evoked by short-timescale input, a complementary component of standardized intrinsic electrophysiological phenotyping^2,4^. We therefore constructed a response-focused brief-pulse geometry from features capturing peak response amplitude, mean response amplitude, fast response magnitude, and input-normalized response gain. This representation isolated fast transient input sensitivity from broader variation in command coverage or stimulation sampling.

The resulting brief-pulse response space formed a structured and low-dimensional geometry **(Fig. 4A-B)**. Cells occupied a continuous response space with biased driver-line occupancy, indicating that transient voltage responses were not distributed randomly across genetically targeted cells. Feature loadings identified the physiological meaning of the leading response axis: response PC1 was dominated by fast and total voltage-response amplitude features, including fast response mean, response mean, response peak, response quantiles, and fast peak response measures **(Fig. 4C)**. This axis captured the magnitude of the rapid voltage response evoked by brief current input, defining a transient-response amplitude geometry.

**Figure 4.**
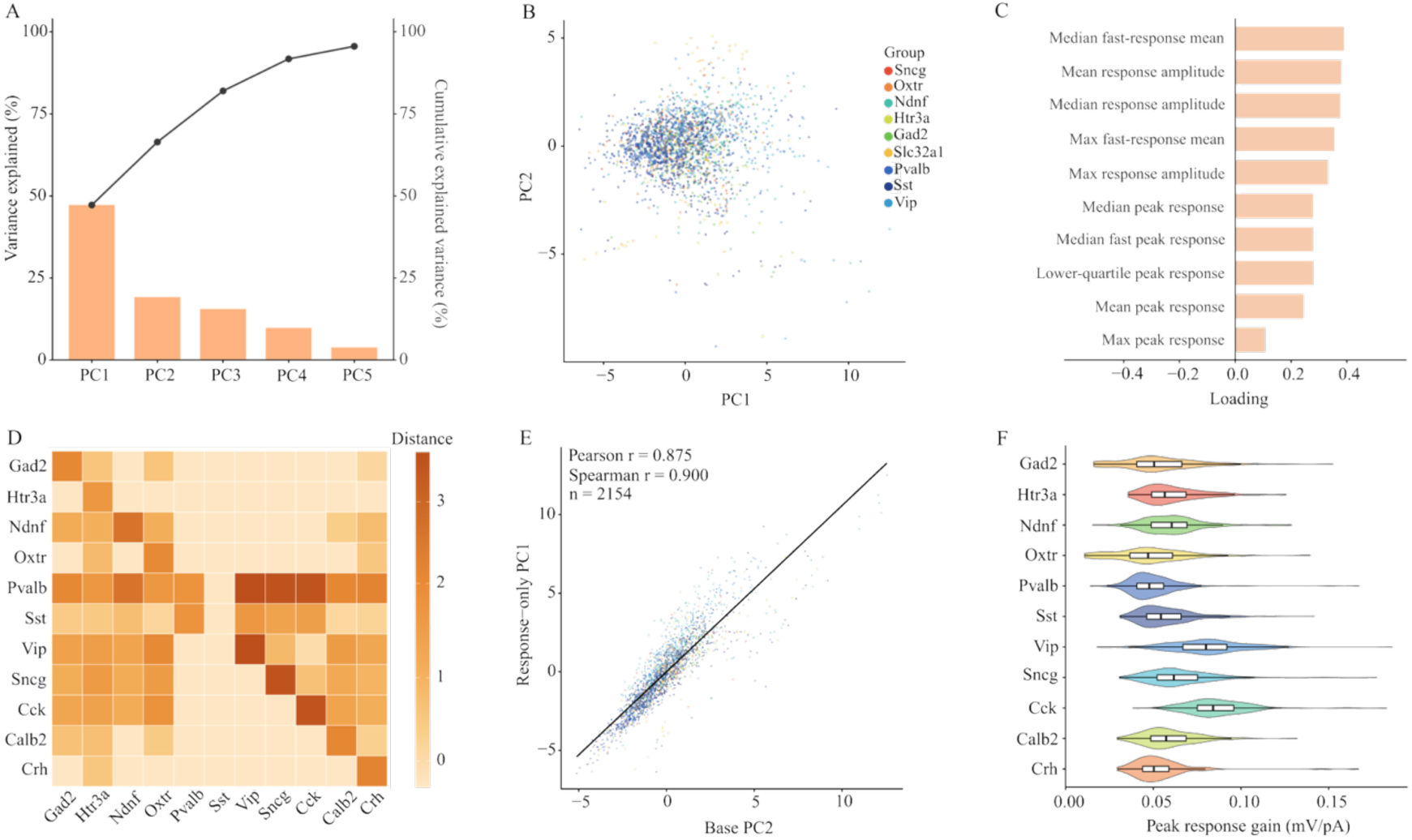
Brief-pulse stimulation defines a fast transient response and gain-related geometry. **(A)** Variance explained by principal components derived from brief-pulse response features. Bars indicate variance explained by individual PCs, and the line indicates cumulative explained variance. **(B)** Cell-level PC1–PC2 embedding of brief-pulse response features, with cells colored by driver-defined group. Driver-line labels are abbreviated by primary marker genes for readability. **(C)** Top response PC1 feature loadings, showing that the leading brief-response axis was dominated by fast and total voltage-response amplitude features. **(D)** Pairwise Euclidean distances between driver-line centroids in brief-pulse response PC space. **(E)** Cell-level alignment between the response-specific brief-pulse geometry and the broader brief-pulse base geometry, showing correspondence between response PC1 and the response-related base PC2 axis. **(F)** Driver-defined group distributions of peak response gain, providing a raw physiological anchor for fast input–response sensitivity.

This transient-response geometry was preserved within the broader brief-pulse feature space. Driver-line centroids were separated in response PC space **(Fig. 4D)**, and response PC1 aligned strongly with the corresponding response-related axis from the broader brief-pulse base geometry **(Fig. 4E)**. This alignment shows that the response-focused representation recovered a major fast-response structure present in the full brief-pulse space, rather than producing a feature-restricted or sampling-specific projection.

Peak response gain provided a raw physiological anchor for this geometry and revealed a driver-group ordering distinct from the long-pulse and ramp protocols **(Fig. 4F)**. Cells with higher gain produced larger voltage responses per unit current input^1,3^, whereas lower-gain cells showed weaker transient sensitivity. Vip- and Cck-targeted cells occupied the higher-gain end of the distribution, whereas Pvalb-targeted cells no longer defined the upper extreme observed in the sustained-output and ramp- recruitment geometries. This reordering demonstrates that fast transient input sensitivity is not simply a scaled version of sustained spiking output or ramp recruitment threshold.

Together, these results establish brief-pulse stimulation as a transient-gain assay that complements and diverges from the long-pulse and ramp geometries. Long-pulse and ramp stimulation organize cells around sustained output and dynamic recruitment, whereas brief-pulse stimulation isolates rapid voltage-response sensitivity. The distinct driver-group ordering in the brief-pulse geometry shows that genetically targeted cell groups express protocol-specific response structure, while still retaining organization within a shared intrinsic electrophysiological landscape.

### Protocol-specific response geometries preserve shared cell-group structure

Long-pulse, ramp, and brief-pulse stimulation each defined a distinct electrophysiological response geometry. We next tested whether these geometries were independent protocol-specific descriptions or different projections of a shared intrinsic organization across driver-defined cell groups, building on prior work showing that cortical cell types can be organized through multivariate electrophysiological and morphoelectric structure^4,5^. This distinction is central: protocol-specific axes show that each stimulation regime emphasizes a different physiological response dimension, but they do not by themselves determine whether cell-group relationships are preserved across regimes.

The strongest axis-level correspondence was observed between long-pulse and ramp stimulation, the two protocols with the greatest cell-level overlap. Driver-line centroids along the long-pulse PC2 output-recruitment axis were closely aligned with centroids along the ramp recruitment PC1 axis **(Fig. 5A)**. This alignment demonstrates that the recruitment hierarchy revealed by ramp stimulation was linked to the sustained-response structure observed under long-pulse stimulation. Thus, long-pulse and ramp protocols captured related aspects of an output-threshold organization across driver-defined groups.

**Figure 5.**
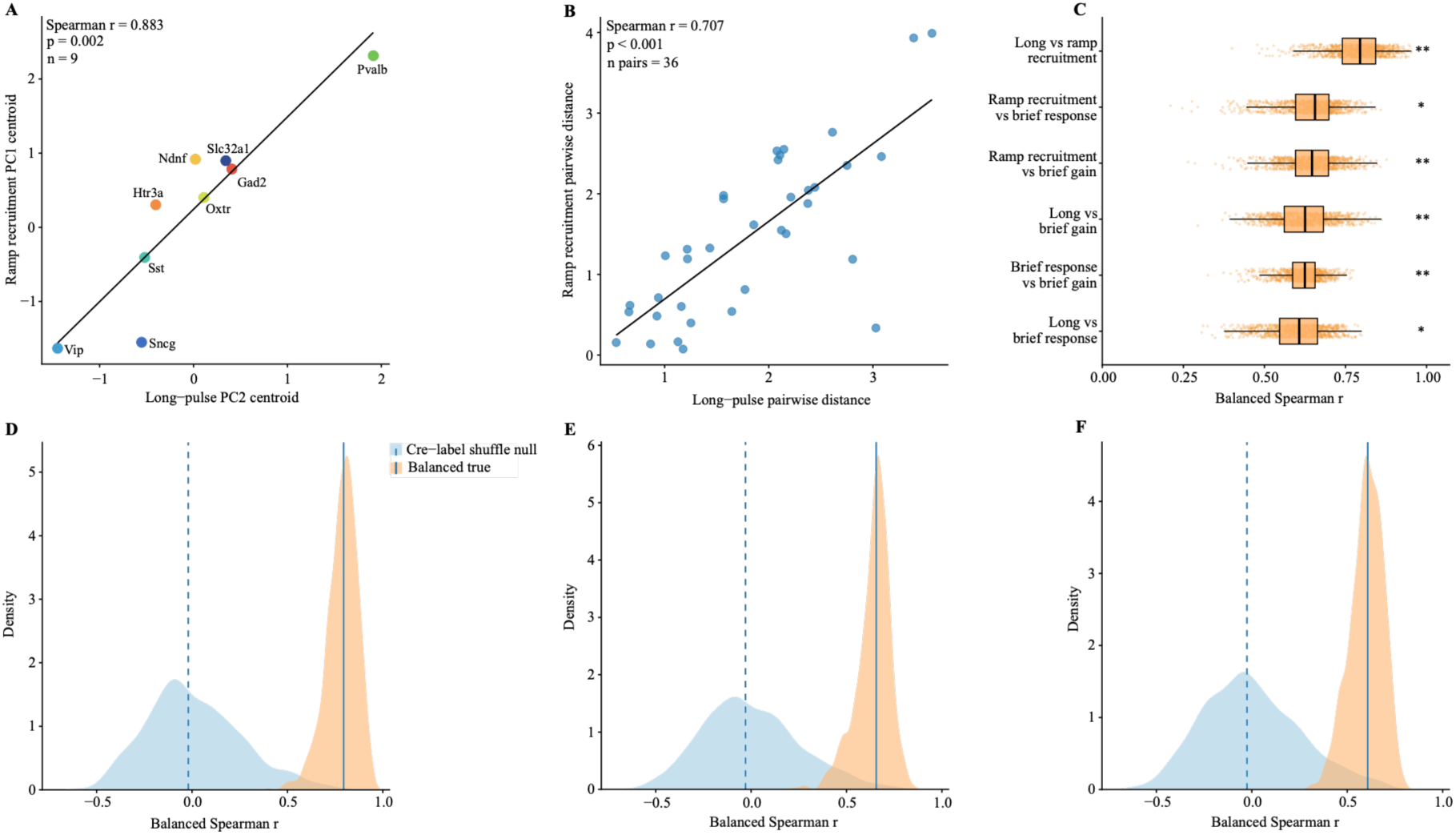
Driver-defined response geometry is preserved across stimulation protocols. **(A)** Driver-line centroid alignment between long-pulse PC2 and ramp recruitment PC1. Each point represents one driver-defined group. **(B)** Pairwise driver-line distance alignment between long-pulse and ramp recruitment geometries. Each point represents one driver-line pair. **(C)** Balanced cross-protocol distance-matrix correlations across protocol comparisons. Boxplots show balanced Spearman correlations across resampling iterations. **(D– F)** Representative balanced validation and driver-label shuffle null distributions for long-pulse versus ramp recruitment, ramp recruitment versus brief-response, and long-pulse versus brief-response comparisons. Orange distributions indicate balanced true alignments, and blue distributions indicate driver-label shuffle null models. True alignments were shifted above the null distributions, indicating that driver-line relational geometry is partially preserved across stimulation regimes.

We then asked whether this correspondence extended beyond a selected pair of axes to the broader relational geometry among cell groups. For each protocol-specific geometry, we computed pairwise distances between driver-line centroids and compared whether driver-line pairs that were far apart in one response space also tended to be far apart in another^11^ **(Fig. 5B)**. Long-pulse and ramp recruitment geometries showed a positive distance-matrix alignment, demonstrating that their relationship was not limited to a single principal component comparison. The relative arrangement of driver-defined groups was preserved across the two response spaces.

To evaluate whether this preservation generalized across stimulation regimes, we extended the distance-matrix analysis to all protocol pairs using balanced resampling **(Fig. 5C-F)**. This procedure matched cell-count structure across driver lines and protocols, allowing cross-protocol geometry to be compared under controlled sampling conditions. Across comparisons, balanced distance-matrix correlations remained consistently positive, including long-pulse versus brief-response, ramp recruitment versus brief-response, ramp recruitment versus brief-gain, long-pulse versus brief-gain, and brief-response versus brief-gain geometries. In each case, the true balanced alignments were shifted above driver-label shuffle null distributions, showing that the preserved geometry could not be explained by unequal sampling or random driver-line assignment.

Together, these analyses establish that driver-defined electrophysiological organization is not confined to a single stimulation protocol. Long-pulse, ramp, and brief-pulse stimulation emphasize different physiological response dimensions—sustained spiking output, dynamic recruitment, and fast transient response/gain—yet preserve a shared relational structure among driver-defined groups. Thus, distinct stimulation protocols do not generate unrelated response spaces. They reveal different physiological projections of a partially shared intrinsic electrophysiological organization. This shared-yet-protocol-specific structure provides a bridge between multivariate single-cell physiology and genetically targeted cell-group identity.

## Discussion

This study establishes a protocol-resolved view of intrinsic electrophysiological organization across genetically targeted cortical neurons. The central aim was not simply to compare driver-defined cell groups under individual stimulation protocols, but to ask whether intrinsic response properties form an organized structure within and across physiological input regimes. Long-pulse, ramp, and brief-pulse stimulation each revealed an interpretable response geometry, and driver-defined groups occupied ordered positions within these spaces. At the same time, cross-protocol comparisons showed that these geometries were not independent descriptions of unrelated response properties. They retained a partially preserved relational structure across stimulation regimes. Together, these findings support a view of electrophysiological identity as a multidimensional response organization, with both protocol-specific expression and shared cell-group structure. This view is consistent with broader efforts to define neuronal identity through combinations of molecular, anatomical, electrophysiological, and functional properties rather than through a single defining feature^12–14^.

The protocol-specific geometries show that intrinsic electrophysiology cannot be reduced to a single excitability scale. Each stimulation protocol imposed a different input demand and therefore emphasized a different transformation from current input to cellular response. Long-pulse stimulation captured how cells convert maintained depolarizing input into sustained spiking output. Ramp stimulation captured the current and timing required to enter the spiking regime during gradually increasing drive. Brief-pulse stimulation captured rapid voltage-response sensitivity and input-normalized gain. These geometries therefore represent complementary response regimes: sustained output, dynamic recruitment, and transient gain. Their separation is important because cells that are prominent along one response dimension need not occupy the same position along another.

Driver-defined group positions illustrate how these response dimensions organize genetically targeted neurons. Pvalb-targeted cells occupied the high end of the long-pulse and ramp-related axes, supporting a high-threshold, high-output intrinsic response mode under sustained and gradually increasing stimulation. They required stronger ramp input for recruitment, yet once sufficiently driven, they occupied the strong-output end of the long-pulse geometry. This profile is consistent with the fast-spiking and high-frequency firing properties commonly associated with Pvalb interneurons, which often contribute to rapid and reliable inhibition in cortical circuits, precise spike timing, and synchronized network activity^15–19^. The interpretation remains grounded in the single-cell recordings: the data identify an intrinsic response regime in which Pvalb-targeted cells transform sufficient input into strong sustained output.

Vip-targeted cells occupied a different region of this response organization. They were positioned lower along the sustained-output and ramp-recruitment axes, but shifted toward the higher-gain end of the brief-pulse geometry. This pattern separates transient voltage-response sensitivity from sustained spiking output and recruitment threshold. VIP interneurons are often enriched in superficial cortical circuits and associated with disinhibitory and state-dependent modulation, including interactions with other inhibitory populations such as Sst interneurons^20–25^. The brief-pulse result therefore provides an intrinsic electrophysiological correlate of a response mode that may be especially sensitive to input timing and circuit state, without treating ex vivo current-clamp responses as direct measurements of circuit function.

The same logic extends beyond the Pvalb/Vip contrast. Cck- and Sncg-targeted cells also contributed to the higher-gain end of the brief-pulse distribution, indicating that transient-response sensitivity is distributed across multiple non-Pvalb inhibitory groups. Electrophysiologically, this axis may reflect variation in membrane filtering, input resistance, capacitance, and fast voltage-response dynamics rather than spike count or recruitment current alone. In cortical inhibitory systems, Cck-, Sncg-, Htr3a-, and Ndnf-related populations are commonly associated with non-Pvalb diversity, including superficial-layer, neurogliaform-like, and modulatory inhibitory circuits^14,16,26–29^. Their positions in the brief-pulse geometry suggest that fast transient response properties provide an additional intrinsic axis for organizing inhibitory diversity beyond sustained output and recruitment threshold.

The continuous structure of the geometries is also informative. Driver-defined groups did not form isolated islands in response space; they occupied biased and partially overlapping regions of shared continua. This pattern is expected for genetic labels that capture heterogeneous biological populations. Sst-targeted cells include dendrite-targeting interneurons with diverse intrinsic and synaptic properties, while broad inhibitory driver lines such as Gad2 and Slc32a1 label large inhibitory classes rather than narrowly defined cell types. Sst and Martinotti-like interneurons have been linked to dendritic inhibition, sensory integration, and context-dependent modulation, but they also include diverse physiological and circuit subtypes^12,30–32^. The observed organization therefore supports a graded view of intrinsic electrophysiological identity. Genetic identity biases cells toward particular regions of physiological response space, but single-cell intrinsic physiology remains continuous within and across driver-defined groups.

Cross-protocol preservation provides the key link between these protocol-specific findings and the broader framework. If long-pulse, ramp, and brief-pulse geometries only reflected unrelated measurements, then group relationships observed under one protocol would not be expected to transfer across stimulation regimes. Instead, distance structures among driver-defined groups remained positively aligned across protocols and exceeded driver-label shuffle null models under balanced resampling. This indicates that protocol-specific response geometries retain a common relational backbone. The strongest correspondence between long-pulse and ramp responses suggests a shared output-threshold organization, while the brief-pulse results show how the same group structure can be re-expressed along a transient-gain dimension. Thus, distinct protocols reveal different physiological projections of a partially shared intrinsic organization.

This view clarifies the contribution of response geometry to electrophysiological phenotyping. Traditional analyses often focus on individual measurements such as firing rate, rheobase, spike threshold, latency, adaptation, input resistance, or response amplitude. These features remain essential, but their interpretation depends on how they covary across stimulation regimes. Response geometry organizes these measurements into structured physiological spaces, allowing both cell-level variation and group-level organization to be examined together. In this framework, electrophysiological identity is not a fixed list of scalar properties. It is an organized response landscape expressed across multiple input regimes, with some dimensions specific to particular protocols and others preserved across protocols. This interpretation is consistent with broader evidence that neuronal classes often combine categorical structure with continuous phenotypic variation across molecular, anatomical, and physiological dimensions^33–36^.

This framework has broader implications for single-cell physiology and cell-type organization. Although the present analysis focused on genetically targeted mouse visual cortical neurons from the Allen Institute Patch-seq dataset, the same logic can be applied to other datasets with standardized stimulation protocols and sufficient cell-group annotation. Other cortical areas, subcortical structures, developmental stages, disease models, perturbation experiments, and species could be represented as protocol-specific response geometries and compared through cell-level embeddings, group centroids, and cross-protocol distance structures. In these settings, biological differences may appear as selective changes in sustained-output, recruitment-threshold, or transient-gain geometries, rather than as uniform shifts in a single electrophysiological feature. The same approach could also be extended beyond driver-line labels to transcriptomic subclasses, morphological types, projection-defined neurons, pharmacological perturbations, or disease-associated cellular states.

Several limitations define the scope of these conclusions. Driver lines are useful genetic entry points but imperfect proxies for biological cell types, and the observed organization should be interpreted as structure among driver-defined groups rather than a complete taxonomy of transcriptomic or circuit-defined cell types. The PCA-based geometries provide interpretable summaries of response organization, but they do not identify the conductances or causal mechanisms that generate the axes. Protocol coverage and same-cell overlap were also incomplete, although balanced resampling and driver-label shuffle controls support the robustness of the cross-protocol alignments. Finally, intrinsic current-clamp responses cannot be directly equated with in vivo circuit function. The known cortical roles of Pvalb, Vip, Sst, Cck, Sncg, Htr3a, and Ndnf populations provide useful context for interpreting group positions, but linking the response modes identified here to circuit computation will require integration with transcriptomic, morphological, anatomical, synaptic, perturbational, and in vivo functional data.

Together, these results show that intrinsic electrophysiological identity is a multidimensional response organization expressed across physiological input regimes. Long-pulse, ramp, and brief-pulse stimulation provide complementary views of sustained output, recruitment threshold, and transient gain, while cross-protocol comparisons reveal partially preserved relational structure among driver-defined groups. Protocol-resolved response geometry therefore provides a framework for connecting multivariate single-cell physiology to biological cell-type organization while preserving the distinction between intrinsic response properties and circuit-level function.

## Methods

### 1. Data source and protocol selection

Electrophysiological recordings were obtained from the publicly available Allen Institute Patch-seq resource, distributed through the DANDI Archive as Dandiset 000020^4,8,37^. We analyzed intracellular current-clamp recordings from genetically defined mouse visual cortical neurons. Raw NWB files were converted into cleaned HDF5 files for downstream analysis. Each cleaned file contained sweep-level voltage responses, current command traces, time vectors, and cell metadata, including driver-line identity. Cre-line labels were used as driver-defined cell-group labels throughout the analysis. Only current-clamp sweeps with successfully loaded voltage, command, and time arrays were retained.

Sweeps were assigned to stimulation protocols using stimulus descriptions, current-command waveform structure, and recording mode. We focused on three stimulation regimes: long-pulse, ramp, and brief-pulse stimulation, corresponding to standardized current-clamp stimulus families used for intrinsic electrophysiological characterization^2,4^. These protocols were selected because they impose distinct input demands on the cell. Long-pulse stimulation measures sustained spiking output under maintained current input. Ramp stimulation measures dynamic recruitment as command current increases over time. Brief-pulse stimulation measures fast transient voltage responses and input-normalized gain.

For the brief-pulse analysis, candidate short-pulse protocols were screened for sweep count, cell coverage, driver-line coverage, pulse detectability, baseline quality, command-amplitude validity, and basic quality-control pass rate. The X5SP_Search_DA_0 protocol was selected as the primary brief-pulse protocol because it provided broad cell and driver-line coverage while maintaining reliable short-pulse detection.

### 2. Sweep-level quality control and feature extraction

Sweep-level quality control and feature extraction were performed separately for each stimulation protocol. Across protocols, quality-control criteria required successful trace loading, current-clamp recording mode, valid trace length, detectable stimulus structure, stable baseline activity, acceptable baseline noise, and successful extraction of response features. Protocol-specific criteria were applied according to stimulus waveform structure: long-pulse sweeps were required to contain a valid sustained current step, ramp sweeps were required to contain a detectable and approximately linear ramp command, and brief-pulse sweeps were required to contain a detectable short current pulse with valid command amplitude and baseline period. Only sweeps passing both common and protocol-specific quality-control criteria were used for feature extraction.

Protocol-native features were extracted from voltage and command traces. Long-pulse features quantified sustained spiking output, voltage displacement, recruitment-related measures, response fractions, input-output slopes, and adaptation-related properties. Ramp features quantified ramp waveform properties, first-spike timing, recruitment current, firing-rate output, spike-count output, and spike acceleration. Brief-pulse features quantified pulse properties, baseline voltage, fast voltage response, response latency, input-normalized gain, spike probability, and first-spike timing when present. A full list of derived features is provided in **Supplementary Table 1**.

### 3. Cell-level aggregation and feature-set definition

Repeated sweeps from the same cell and protocol were aggregated into one cell-level feature vector. Aggregation used robust summary statistics, including medians, means, minima, maxima, quantiles, response fractions, spike fractions, and input-output slopes where appropriate. This produced separate cell-level feature tables for long-pulse, ramp, and brief-pulse stimulation.

For each downstream geometry, features were selected according to the physiological regime being analyzed. Long-pulse analyses used a sustained-response feature set. Ramp analyses included a broad ramp feature set and a recruitment-focused feature set; the recruitment-focused geometry was used as the main ramp representation for cross-protocol comparison. Brief-pulse analyses included response-focused and gain-focused feature sets, allowing fast transient voltage-response amplitude to be separated from input-normalized gain.

### 4. Protocol-specific response geometries

For each protocol-specific feature matrix, features were filtered for excessive missingness, missing values were imputed, and retained features were standardized before dimensionality reduction.

Principal component analysis was performed separately for each protocol and feature set. This generated protocol-specific electrophysiological response geometries in which each cell was represented by its scores along principal component axes.

For each PCA geometry, we exported cell PC scores, feature loadings, explained variance ratios, driver-line centroids, and pairwise driver-line centroid distances. Feature loadings were used to interpret the physiological meaning of major axes, and explained variance ratios were used to quantify how much response variability was captured by leading components. Because PCA signs are arbitrary, interpretation focused on feature composition, relative cell positions, and cross-geometry alignment rather than the absolute sign of any principal component.

Long-pulse PCA was used to define a sustained spiking-output geometry. Ramp PCA was used to define a dynamic recruitment geometry, with the recruitment-focused feature set used for the main ramp analysis. Brief-pulse PCA was used to define response-focused and gain-focused geometries, capturing fast transient voltage-response amplitude and input-normalized gain, respectively.

### 5. Driver-line centroids and cross-protocol distance alignment

Within each protocol-specific PCA space, driver-line centroids were calculated as the mean PC score vector of cells belonging to each driver line. Pairwise Euclidean distances between driver-line centroids were computed to form a driver-line distance matrix for each protocol geometry. These distance matrices served as the main representation of driver-line relational geometry.

Cross-protocol alignment was quantified by comparing driver-line distance matrices across protocol-specific geometries. For each pair of geometries, shared driver lines were identified, and the upper triangle of each pairwise centroid distance matrix was vectorized. Pearson and Spearman correlations were then computed between distance vectors. This analysis tested whether driver-line pairs that were far apart in one protocol geometry also tended to be far apart in another, even when the raw features used to construct the geometries differed across protocols.

For long-pulse and ramp geometries, same-cell comparisons were also performed because many cells were represented in both protocols. Cells were matched by identity, and correlations were computed between long-pulse PC scores, ramp PC scores, and raw ramp recruitment features. For comparisons involving brief-pulse geometry, analyses were performed at the driver-line centroid and distance-matrix levels because the brief-pulse cell set had limited overlap with the long-pulse and ramp cell sets.

### 6. Balanced resampling and label-shuffle null models

Balanced resampling was used to control for unequal driver-line sample sizes across protocols. For each cross-protocol comparison, driver lines with insufficient representation across the required protocol geometries were excluded. In each iteration, the same number of cells was randomly sampled from each retained driver line within each protocol geometry. Driver-line centroids and pairwise distance matrices were recomputed from the balanced samples, and cross-protocol distance-matrix correlations were recalculated.

Label-shuffle null models were generated by permuting driver-line labels within each balanced sample while preserving the PC score distribution and sample-size structure. Centroids, distance matrices, and cross-protocol correlations were recomputed after label shuffling. Empirical p-values were calculated by comparing balanced true correlations against the corresponding shuffled-label null distributions. This procedure tested whether cross-protocol alignment persisted after controlling for driver-line sample imbalance and whether the observed alignment depended on true driver-line identity.

### 7. Implementation and output files

All analyses were performed using custom Python scripts. HDF5 files were accessed with h5py; tabular data were processed with pandas and NumPy; scaling, imputation, and PCA were performed with scikit-learn; statistical correlations and resampling analyses were performed using standard scientific Python libraries; and plots were generated with matplotlib^38–41^.

Intermediate and final outputs were saved as CSV and JSON files, including quality-control summaries, protocol-specific feature tables, PCA scores, feature loadings, explained variance tables, driver-line centroids, centroid distance matrices, cross-protocol correlations, balanced resampling results, and label-shuffle null distributions.

## Acknowledgments

This study utilized publicly available datasets from the Allen Brain Observatory, provided by the Allen Institute for Brain Science. The authors gratefully acknowledge funding from the Westlake Fellows Program at Westlake University.

## Author contributions

Conceptualization: Z.Y., Y.X. Methodology: Z.Y. Software: Z.Y. Formal analysis: Z.Y. Investigation:

Z.Y. Visualization: Z.Y. Writing – original draft: Z.Y. Writing – review & editing: Z.Y., Y.X. Supervision: Y.X. Project administration: Y.X.

## Data and Code Availability

The electrophysiological recordings analyzed in this study are publicly available from the Allen Institute for Brain Science Patch-seq dataset hosted on the DANDI Archive as Dandiset 000020 (DOI: 10.48324/dandi.000020/0.210913.1639). The analysis code used in this study will be made publicly available upon publication.

