## Supplementary Table 1 for "Electrophysiological response geometries reveal shared and protocol-specific cell-type organization"

**Supplementary Table 1. Protocol-specific derived features used for response geometry construction.** Derived electrophysiological features are grouped by stimulation protocol and feature class. Long-pulse features summarize sustained spiking output, voltage-response amplitude, command-binned responses, and baseline properties. Ramp features summarize recruitment current, first-spike timing, post-recruitment output, voltage-response amplitude, and baseline properties. Brief-pulse features summarize command coverage, transient voltage-response amplitude, input-normalized gain, response timing, spike probability, and baseline properties. Feature classes were used to construct protocol-specific cell-level feature matrices for PCA-based response geometry analysis.

**Long-pulse features**

| Feature class | Derived features |
|---|---|
| Response fractions | Spiking fraction; subthreshold-response fraction; strong-depolarization fraction; high-spiking fraction |
| Sustained firing output | Median firing rate; mean firing rate; maximum firing rate |
| Spike-count output | Median spike count; maximum spike count |
| Recruitment-related response | First-spiking command amplitude |
| Input-output relationship | Firing-rate slope; peak-response slope |
| Voltage-response amplitude | Median peak response amplitude |
| Baseline properties | Median baseline membrane potential; median baseline noise |
| Command-binned response fractions | Spiking fractions across depolarizing command-amplitude bins |
| Command-binned voltage responses | Median peak response amplitudes across depolarizing command-amplitude bins |
| Command-binned firing output | Median firing rates across depolarizing command-amplitude bins |

**Ramp features**

| Feature class | Derived features |
|---|---|
| Recruitment current | Median recruitment current; minimum recruitment current |
| Spiking probability | Spiking fraction |
| First-spike timing | Median first-spike latency; minimum first-spike latency |
| Spike-count output | Median spike count; maximum spike count |
| Post-recruitment firing output | Median firing rate; maximum firing rate |
| Voltage-response amplitude | Median peak response amplitude; median mean response amplitude; median absolute peak voltage |
| Baseline properties | Median baseline membrane potential; median baseline noise |

**Brief-pulse features**

| Feature class | Derived features |
|---|---|
| Protocol coverage | Brief-pulse sweep count; unique command-level count |
| Command properties | Median command amplitude; maximum command amplitude; command-amplitude range |
| Baseline properties | Median baseline membrane potential; median baseline noise |
| Peak voltage response | Median peak response amplitude; maximum peak response amplitude |
| Mean voltage response | Median mean response amplitude |
| Fast voltage response | Median fast-response peak amplitude; maximum fast-response peak amplitude; median fast-response mean amplitude |
| Input-normalized gain | Median peak-response gain; median mean-response gain |
| Command-response relationship | Peak-response command slope; peak-response slope fit |
| Response timing | Median peak-response latency; median fast-response peak latency |
| Spike probability and output | Spike fraction; maximum spike count |
| First-spike recruitment and timing | First-spiking command amplitude; median first-spike latency |
